# Arterial endothelial deletion of hereditary hemorrhagic telangiectasia 2/*Alk1* causes epistaxis and cerebral microhemorrhage with aberrant arteries and defective smooth muscle coverage

**DOI:** 10.1101/2024.11.25.622742

**Authors:** Xuetao Zhang, Kyle A Jacobs, Kunal P Raygor, Shang Li, Jiajun Li, Rong A Wang

## Abstract

Hereditary Hemorrhagic Telangiectasia (HHT) is an autosomal dominant vascular disorder with manifestations including severe nose bleeding and microhemorrhage in brains. Despite being the second most common inherited bleeding disorder, the pathophysiological mechanism underlying HHT-associated hemorrhage is poorly understood. HHT pathogenesis is thought to follow a Knudsonian two-hit model, requiring a second somatic mutation for lesion formation. Mutations in activin receptor-like kinase 1 (*ALK1*) gene cause HHT type 2. We hypothesize that somatic mutation of *Alk1* in arterial endothelial cells (AECs) leads to arterial defects and hemorrhage. Here, we mutated *Alk1* in AECs in postnatal mice using *Bmx(PAC)-Cre^ERT2^* and found that somatic arterial endothelial mutation of *Alk1* was sufficient to induce spontaneous epistaxis and multifocal cerebral microhemorrhage. This bleeding occurred in the presence of tortuous and enlarged blood vessels, loss of arterial molecular marker *Efnb2*, disorganization of vascular smooth muscle, and impaired vasoregulation. Our data suggest that arterial endothelial deletion of *Alk1* leading to reduced arterial identity and disrupted vascular smooth muscle cell coverage is a plausible molecular mechanism for HHT-associated severe epistaxis. This work provides the first evidence that somatic *Alk1* mutation in AECs can cause hemorrhagic vascular lesions, offering a novel preclinical model critically needed for studying HHT-associated epistaxis, and delineating an arterial mechanism to HHT pathophysiology.

## Introduction

Hereditary hemorrhagic telangiectasia (HHT), also known as Osler–Weber–Rendu syndrome, is an autosomal dominant vascular disorder characterized by bleeding and malformations of blood vessels^1,2^. One of the most common clinical presentations, and one of the major diagnostic criteria^1^, is severe recurring epistaxis, or nose bleeding, which was reported to occur in up to 96% of HHT patients^3^. Severe bleeding frequently leads to secondary complications, such as anemia, in up to 50% of HHT patients^4^. HHT patients additionally present with multifocal arteriovenous malformations (AVMs), blood vessels that shunt arterial blood directly into veins without passing through the capillaries^5,6^, of the brain^7^ and other tissues^2^. These peripheral vascular abnormalities can result in subsequent development of high-output cardiac failure which increases the risk of HHT-associated mortality^8^. Despite these consequential impacts on HHT patient quality of life, there are no FDA- or EMA-approved therapeutics or prophylactics for HHT-associated bleeding and patients often require surgical intervention to address bleeding lesions.

The pathological basis of HHT-associated epistaxis is not well understood. Bleeding in HHT patients typically originates from Little’s area, also known as Kiesselbach’s plexus, a vascular plexus on the anterior nasal septum formed by the anastomosis of the anterior and posterior ethmoidal arteries, the sphenopalatine artery, the greater palatine artery, and the superior labial artery^9,10^. The nasal vasculature of children with a parent affected by HHT exhibits both focal telangiectasias and varicose vessels^11^. The data indicate that in some patients, no telangiectasias were detected in the nose, however the patient nonetheless developed recurrent epistaxis^11^. These clinical findings of epistaxis with undetectable telangiectasia raise question of other vascular causes of epistaxis. It is also unclear whether severity of epistasis correlate to the size of nasal telangiectasia. Furthermore, defects in arterial flow are likely to explain severe bleeding. Therefore, it is a reasonable to question whether lesions in feeding artery may contribute to HHT-associated epistaxis.

HHT type-2 (HHT2) is a subtype of HHT characterized by germline inheritance of a mutated copy of *ALK1/HHT2*^12–14^. ALK1, or activin receptor-like kinase 1, is a receptor tyrosine kinase in the TGFβ/BMP pathway^15^. Mouse models have been instrumental in understanding the genetic underpinnings of HHT-associated vascular malformations^16,17^. While germline loss of *Alk1* is embryonic lethal by E10.5 in mice^13^, *Alk1^+/-^* mice are fertile and viable and may develop vascular malformations with low frequency and latency^18^. Studies examining the effects of somatic *Alk1* mutation utilizing tissue-specific mouse genetic strategies have been facilitated by the generation of an *Alk1-*floxed allele^19^. Interestingly, homozygous deletion of *Alk1* mediated by *L1cre,* a transgenic *Alk1* promoter-driven Cre allele, resulted in AVMs and embryonic lethality by E18.5^19^. *Alk1* deletion using *Rosa26Cre^ER^*, an ubiquitous Cre allele, shows that global *Alk1* loss postnatally causes de novo AVMs in response to tissue injury and subsequent moribundity^20^. Postnatal deletion of *Alk1* specifically in endothelial cells, utilizing a pan endothelial Cre allele, *Cdh5(PAC)-Cre^ERT2^*, induces the formation of retinal AVMs^21,22^. Despite these progresses, HHT-associated epistaxis has not been reported in mouse models. Thus, there is a critical need to uncover the pathogenic mechanisms leading to vascular hemorrhage to inform the development of novel HHT therapeutics and prophylactics.

With an autosomal dominant pattern of inheritance, all cells in the body lack a functional copy of *ALK1/HHT2*. However, vascular malformations in HHT patients appear focally and stochastically. This dichotomy has led the hypothesis that HHT-associated vascular malformations may follow a Knudsonian, two-hit model of inheritance, whereby a second somatic mutation is required for the development of a vascular malformation^23,24^. Recent work has utilized next generation sequencing to investigate loss of heterozygosity events in HHT AVMs and identified somatic mutations which support this bi-allelic two-hit mutation model^25,26^. Since this second somatic mutation may occur in any cell of the vasculature, these findings raise pressing questions as to whether the site of the somatic mutation may impact phenotypic presentation. Previous work has elegantly demonstrated that loss of HHT genes in venous and capillary endothelial cells leads to the development of AVMs in mice and zebrafish^27–31^. Here we investigate the arterial contribution to HHT pathogenesis and ask whether somatic mutation of *Alk1* in the arterial endothelial cells (AECs) is sufficient to induce formation of HHT-associated epistaxis and vascular malformations.

To examine the role of arterial endothelial *Alk1* in driving HHT pathogenesis, we utilized the *Bmx(PAC)-Cre^ERT2^* allele^27,32^ to induce mutation of *Alk1* in primarily AECs of postnatal mice. Excitingly, somatic mutation of *Alk1* in AECs caused spontaneous epistaxis and cerebrovascular hemorrhage, malformed arteries with defective arterial specification, smooth muscle coverage, and vasoregulation. This new-found appreciation of the arterial contribution to HHT pathogenesis in mice may inspire future studies into arterial contribution to bleeding in HHT patients.

## Results

### Somatic loss of ALK1 in cells of the *Bmx(PAC)-Cre^ERT2^* lineage induced epistaxis and cerebral microhemorrhage

To determine the effects of loss of *Alk1* in AECs in mice, we somatically mutated both *Alk1* alleles in the cells of the *Bmx(PAC)-Cre^ERT2^* lineage, which consists primarily of AECs^27,32,33^. We activated *Bmx(PAC)-Cre^ERT2^* activity in mice carrying both *Alk1* alleles flanked by loxP sites [*Bmx(PAC)-Cre^ERT2^*; *Alk1^fx/fx^*, hereafter *Alk1^i^*^Δ*AEC*^]. As controls, we used mice carrying *Alk1^fx/fx^* without the *Bmx(PAC)-Cre^ERT2^* allele (hereafter control).

To evaluate the *Bmx(PAC)-Cre^ERT2^* recombinase-mediated depletion of ALK1, we conducted wholemount immunostaining for ALK1 in the cerebral cortical slices of postnatal day (P) 31 *Alk1^i^*^Δ*AEC*^ and control mice injected with 25µg tamoxifen on P2 (**Fig. 1A**). We found that *Alk1^i^*^Δ*AEC*^ mice exhibited depletion of ALK1 in the arteries but retained ALK1 in the veins and capillaries. We also performed ALK1 immunostaining in the cerebral cortical slices of P21 *Alk1^i^*^Δ*AEC*^ and control mice injected with 100µg tamoxifen on P2 and P3 and show similar depletion of ALK1 in the arteries but retained in the veins and capillaries (**Supplemental Fig. 1**). To further confirm the arterial specificity of ALK1 depletion, we used *Alk1^i^*^Δ*AEC*^ and control mice also carrying the *Efnb2^H2B-GFP^* arterial endothelial reporter^34^. This allele expresses Histone H2B-GFP under the endogenous *Efnb2* promoter, thereby labeling the nuclei of arterial, but not capillary or venous, endothelial cells^34^. Indeed, we find that ALK1 depletion is restricted to vessels with nuclear GFP signal, or AECs (**Fig. 1B**). Taken together, these data demonstrate the arterial endothelial specificity of ALK1 protein depletion in *Alk1^i^*^Δ*AEC*^ mice.

**Fig. 1.**
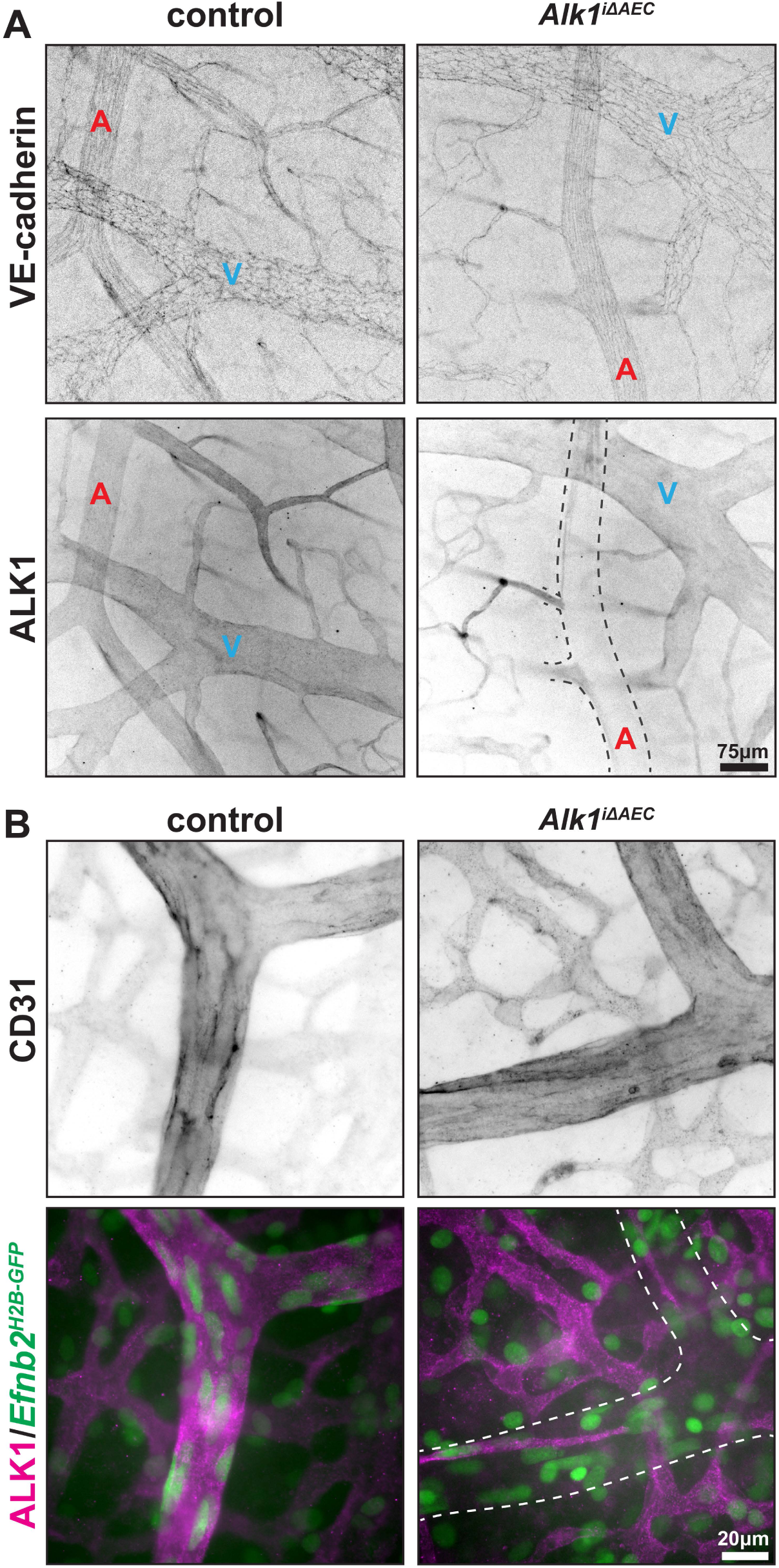
*Bmx(PAC)-Cre^ERT2^* mediated depletion of ALK1 was restricted to arterial endothelium. **(A)** Wholemount immunostaining of cerebral cortex slices for VE-cadherin and ALK1 from P31 control and *Alk1^iΔAEC^* mice (25µg tamoxifen on P2). **(B)** Wholemount immunostaining for CD31 and ALK1 in cerebral cortex slices of P10 control and *Alk1^iΔAEC^* mice expressing *Efnb2^H2B-GFP^* (50µg tamoxifen on P2). A and V label arteries and veins, respectively. Dashed outlines in ALK1 images indicate artery position.

*Alk1^i^*^Δ*AEC*^ mice exhibited spontaneous episodes of epistaxis from about 4 weeks of age (**Fig. 2A**). Epistaxis episodes were identified during daily health checks wherein blood was found dripped on bedding or sprayed on cage walls (**Fig. 2B**). We documented epistaxis or blood on bedding/cage walls in 47.6% of cages containing *Alk1^i^*^Δ*AEC*^ mice (N=21 cages, both male and female mice). We also randomly identified mice with blood on the tips of their noses (**Fig. 2A**). Fortuitously, we found that upon holding the mice for examination, epistaxis would frequently occur (**Supplemental Movie 1**), confirming the nose as the source of bleeding. These epistaxis events were transient in nature with the bleeding quickly stopping and mice rapidly cleaning the blood off the nose. With this transient bleeding, *Alk1^i^*^Δ*AEC*^ mice lived and continued exhibiting epistaxis as long as we kept them (longest is P195), resembling HHT patients with nosebleeds.

**Fig. 2.**
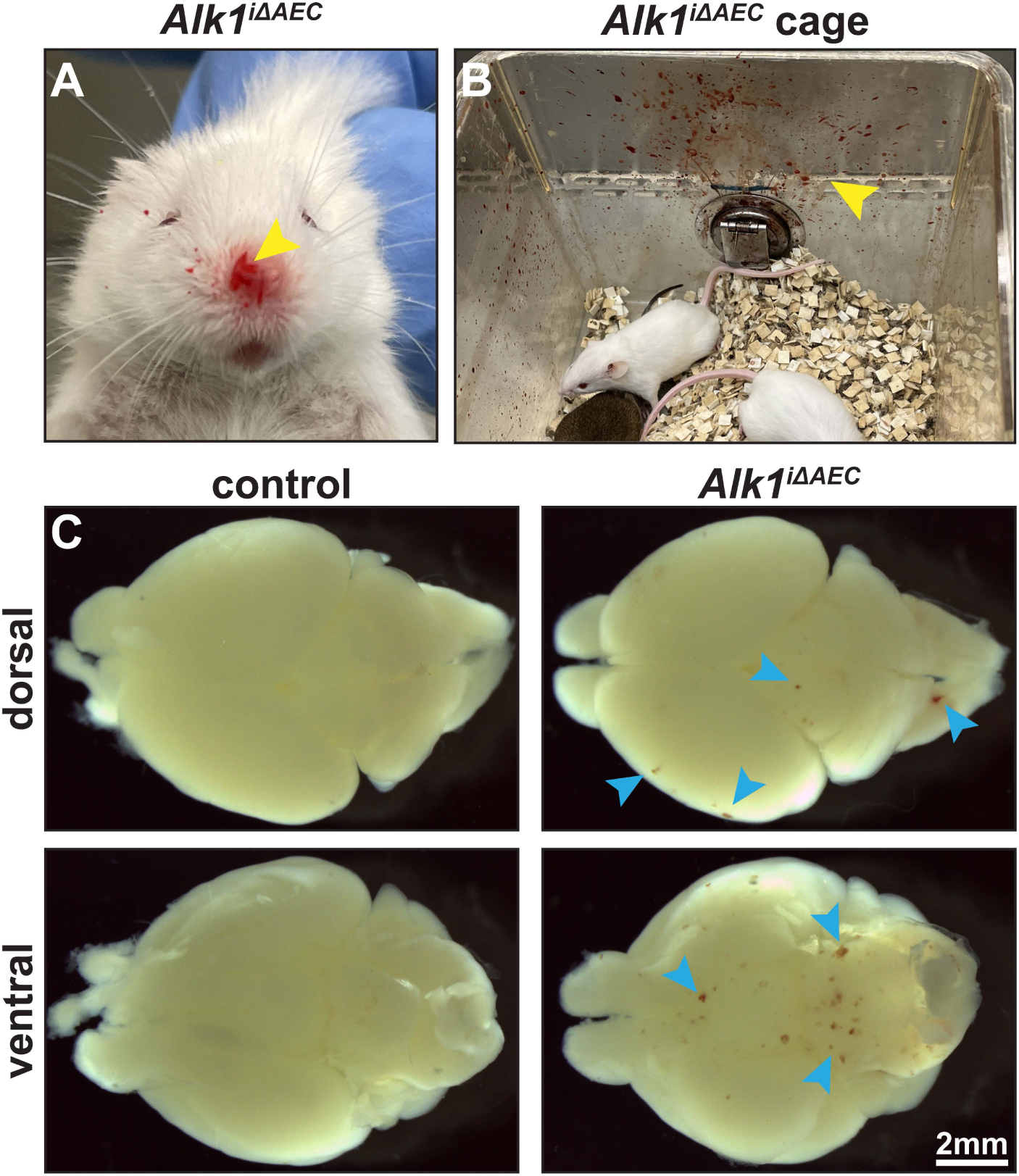
*Alk1^iΔAEC^* mice exhibited epistaxis and brain hemorrhage. **(A)** Photograph of P33 *Alk1^iΔAEC^* mouse with blood on nose (arrowhead). **(B)** Photograph of blood present (arrowhead) in cage housing P81 *Alk1^iΔAEC^* mice. **(C)** Gross pathological images demonstrating multi-focal hemorrhages (arrowheads) present in *Alk1^iΔAEC^* mice brains but absent in control at P7. All mice administered 25µg tamoxifen on P2.

To determine the brain phenotypes in *Alk1^i^*^Δ*AEC*^ mice, we conducted necropsies and found that 72.4% of *Alk1^i^*^Δ*AEC*^ mice displayed multifocal microhemorrhages located throughout the brain (N=58 mice; **Fig. 2C**). We documented no significant difference in incidence cerebral vascular hemorrhage in male and female *Alk1^i^*^Δ*AEC*^ mice. Thus, arterial endothelial loss of *Alk1* is sufficient for the development of multifocal microhemorrhages. As patients with HHT2 also develop multifocal microhemorrhages in the brain^35^, *Alk1^i^*^Δ*AEC*^ mice closely resembled this phenotypic presentation.

### A viable mouse model of HHT2 with epistaxis and cerebral microhemorrhage exhibited dermal telangiectasias and compensatory cardiac failure

In optimizing this preclinical model, we aimed to identify permissive conditions in which mice developed epistaxis and other vascular abnormalities without developing secondary systemic comorbidities that lead to declined health and moribundity shortly after *Alk1* mutation. To this end, we tested different tamoxifen doses for Cre recombinase induction (25, 50, 75, and 100µg). We found that at high dose (100µg on P2 and P3), *Alk1^i^*^Δ*AEC*^ mice developed malformations described here with quick onset and more uniform phenotypes at a given time, however, rapidly became moribund (median 23 days; **Fig. 3**). Low dosing of tamoxifen (25µg) led to slower onset of and milder phenotype, still sufficient to induce vascular malformations. Low tamoxifen doses very rarely caused systemic illness. We likewise administered 50µg and 75µg tamoxifen on P2; these doses resulted in the same vascular phenotypes as 25µg but with sooner onset and variable severity. Together, this dosing strategy provides the framework for an HHT model with tamoxifen-titratable phenotypic severity. We thus primarily utilized 25µg tamoxifen on P2 or 100µg tamoxifen on P2 and P3 in our investigations.

**Fig. 3.**
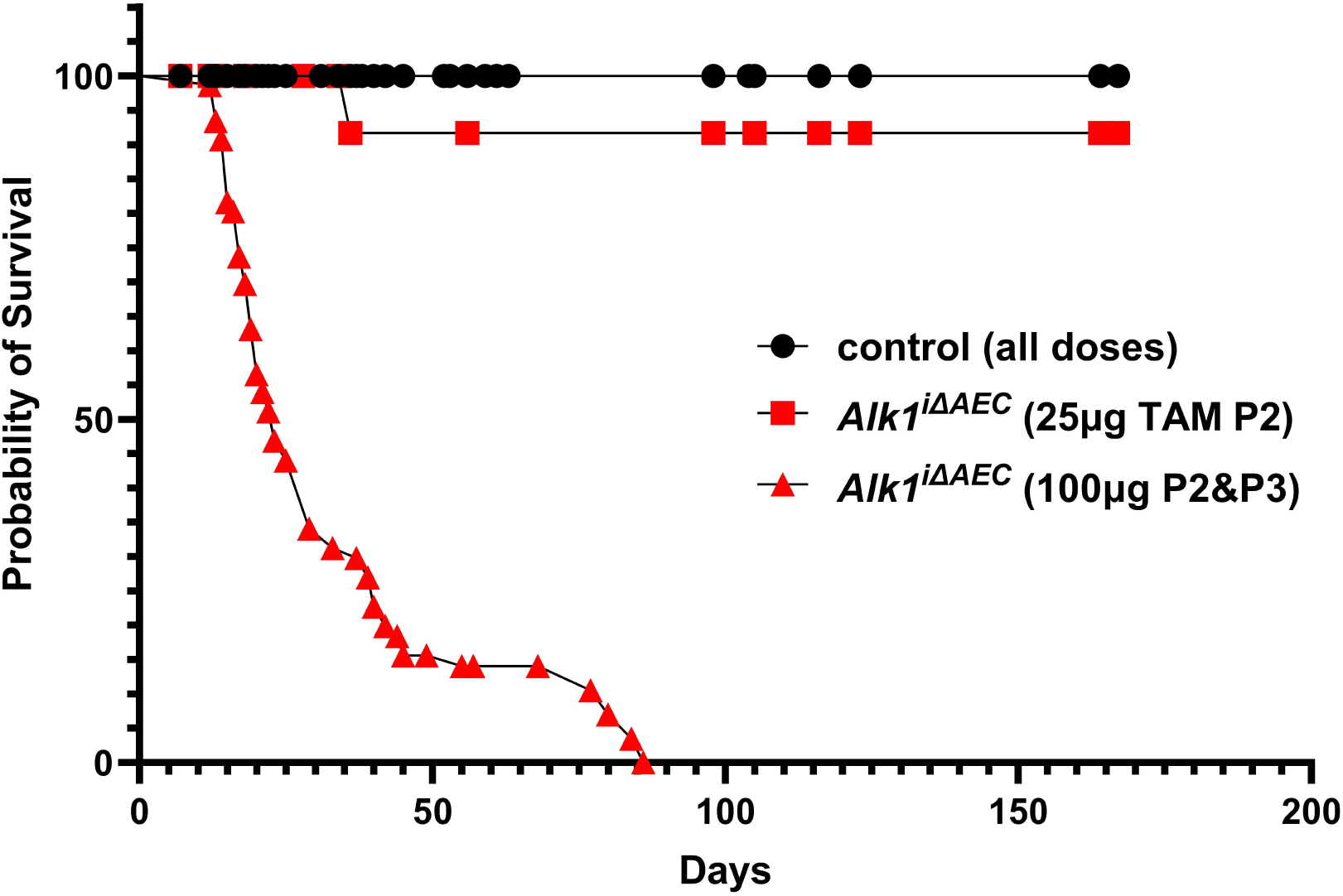
*Alk1^iΔAEC^* mice exhibited tamoxifen dose-dependent moribundity. Kaplan-Meier analysis showed that time to moribundity in *Alk1^iΔAEC^* mice administered 100µg tamoxifen on P2 and P3 (red triangles) was significantly faster (median 23 days) than control (black circles) and *Alk1^iΔAEC^* mice administered 25µg tamoxifen on P2 (N = 45-80 mice per condition).

HHT patients can develop high-output heart failure due to peripheral vascular shunting^36^. To evaluate cardiomegaly, an indicator of high-output cardiac failure, we documented heart to body weight ratio, which reflects cardiomegaly. We found that *Alk1^i^*^Δ*AEC*^ mice developed an enlarged heart and a greater heart weight to body weight ratio relative to controls, over two weeks of age, when tamoxifen is administered at P2 (**Supplemental Fig. 2**). Because *Alk1^i^*^Δ*AEC*^ mice develop brain hemorrhages as early as P7 when tamoxifen is administered at P2 (**Fig. 2C**), we also examined mice between P7 and P15, to determine if a change in heart size succeeds vascular phenotypes. We found no significant difference in heart weight to body weight ratio in mice younger than 15 days old (**Supplemental Fig. 2**), indicating that changes in vascular phenotype preceded compensatory cardiomegaly development. The development of compensatory high-output cardiac failure in *Alk1^i^*^Δ*AEC*^ mice demonstrates further that arterial endothelial loss of ALK1 recapitulates this clinical presentation observed in HHT patients.

We also observed the formation of dermal vascular malformations in *Alk1^i^*^Δ*AEC*^ mice, like seen in HHT patients^37^. These vascular malformations were most evident on the dorsal head skin of *Alk1^i^*^Δ*AEC*^ mice receiving higher doses of tamoxifen (**Supplemental Fig. 3**) and presented heterogeneously, with some being focal, and others being diffuse tangles of tortuous vessels. Thus, administering *Alk1^i^*^Δ*AEC*^ mice higher doses of tamoxifen leads to dermal vascular malformations, further validating the model’s phenotypic similarity to human.

To address whether deletion at later timepoint is sufficient to induce phenotypic presentation, we administered 100µg tamoxifen per g body weight, or high dose, at P13 and P14 and sacrificed mice at 14 weeks of age. We found these mice exhibited epistaxis (**Supplemental Fig. 4A**) and cardiomegaly with average heart weight to body weight ratio of *Alk1^i^*^Δ*AEC*^ and control mice being 0.010 and 0.0045, respectively (N=2 *Alk1^i^*^Δ*AEC*^ mice; **Supplemental Fig. 4B**).

In summary, we show that Cre recombinase induction by injection of 25µg tamoxifen on P2 leads to a progressive phenotypic development gradually with ongoing transient epistaxis, while higher dose leads to a quicker phenotypic onset and more uniform phenotypes at a given time.

### *Alk1^i^*^Δ^*^AEC^* mice exhibited blood vessel enlargement and tortuosity

Given the spontaneous bleeding in *Alk1^i^*^Δ*AEC*^ mice, we sought to determine gross changes in the nasal vasculature contributing to the development of hemorrhage. It is well established that most human epistaxis is caused by hemorrhage of the vasculature of Little’s area/Kiesselbach’s plexus^9,10^. To visualize the feeding arteries and draining veins of the murine structure analogous to Little’s area we casted *Alk1^i^*^Δ*AEC*^ and control mice using MICROFIL and optically cleared the head to visualize blood vessels on the cranial exterior. We discovered a significant enlargement of the facial artery/vein and their tributaries in the mutant mice. We furthermore identified a bed of tortuous blood vessels in the distal nasal plate, which was not detectable in control mice (**Fig. 4A**). Thus, by vascular casting we demonstrate significant vascular enlargement and tortuosity in the head of *Alk1^i^*^Δ*AEC*^ mice.

**Fig. 4.**
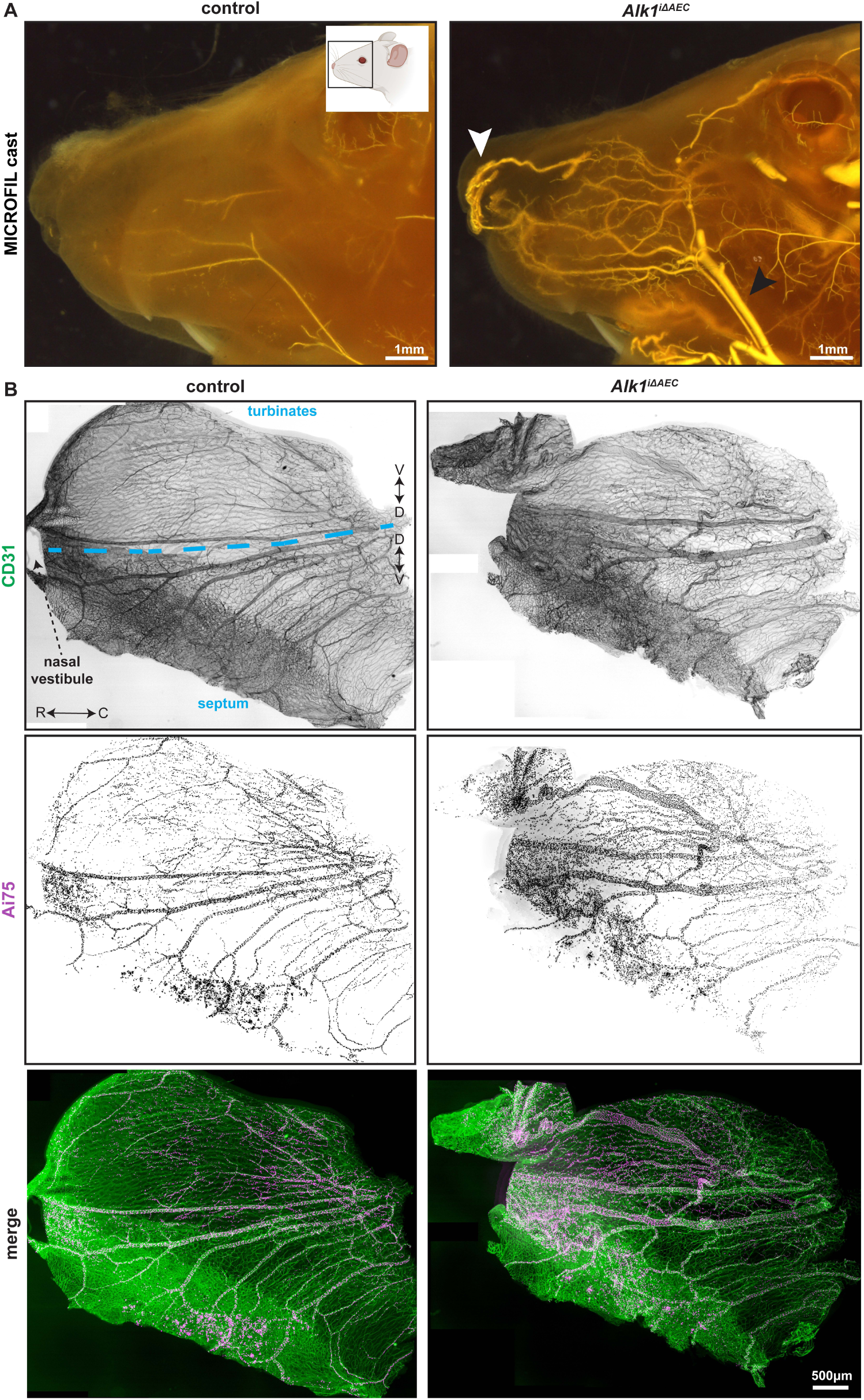
*Alk1^iΔAEC^* mice exhibited enlarged and tortuous nasal vasculature. **(A)** MICROFIL casting of nasal cavity and cranial exterior of control (P16) and *Alk1^iΔAEC^* (P19) mice revealed enlarged and tortuous vessels in *Alk1^iΔAEC^* mouse (100µg tamoxifen on P2 & P3). Black and white arrowheads indicate the facial artery/vein pair and a tangle of blood vessels in the distal nasal vestibule, respectively. Mouse schematic created in BioRender.com. **(B)** Wholemount immunostaining for CD31 and *Rosa26^Ai75^* Cre reporter signal in nasal mucosa of P21 control (*Bmx-Cre^ERT2^*; *Alk1^fx/+^*) and *Alk1^iΔAEC^* mice (25µg tamoxifen on P2). Blue dashed line indicates the most dorsal crease of the nasal cavity. Caudal (C), rostral (R), dorsal (D), and ventral (V) directions are indicated.

To further investigate the vascular defects within the nasal cavity where bleeding was occurring, we conducted wholemount immunofluorescent staining of the nasal mucosa of *Alk1^i^*^Δ*AEC*^ and control mice which revealed enlarged and tortuous large vessels within the nasal mucosa. Interestingly, the vessels exhibiting enlargement were arteries which had undergone Cre-mediated recombination and loss of *Alk1*, as demonstrated by expression of the *Rosa26^Ai75^* Cre recombinase reporter (**Fig. 4B**). Taken together, these results indicate that *Alk1^i^*^Δ*AEC*^ mice exhibited enlarged and tortuous major vessels.

We likewise interrogated the vascular patterning of the cerebral cortex where multifocal hemorrhages formed in *Alk1^i^*^Δ*AEC*^ mice. To visualize the vasculature of the brains of *Alk1^i^*^Δ*AEC*^ and control mice, we conducted MICROFIL casting which revealed that the vessels of *Alk1^i^*^Δ*AEC*^ mice are significantly more tortuous than those of the control (**Fig. 5A**). To investigate this phenotype in higher resolution using a complimentary modality, we immunostained the cerebral cortex of *Alk1^i^*^Δ*AEC*^ and control mice for CD31 and confirmed that the large vessels exhibit tortuosity (**Fig. 5B**). Thus, we find that arterial endothelial mutation of *Alk1* is sufficient to drive the development of large vessel tortuosity in the brain.

**Fig. 5.**
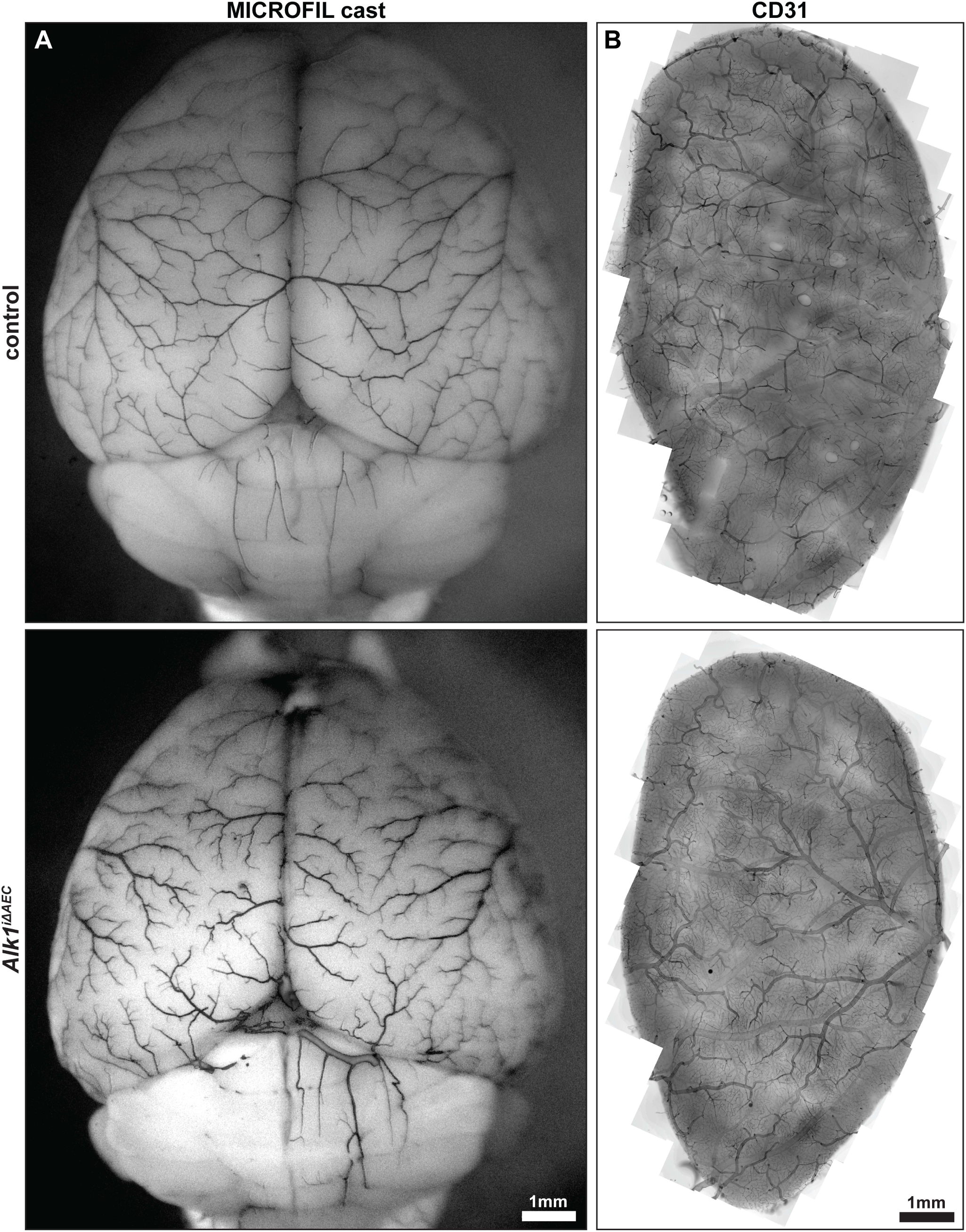
*Alk1^iΔAEC^* mice exhibited tortuous cerebral vasculature. **(A)** MICROFIL casting of the brain demonstrating tortuous vessels in control and *Alk1^iΔAEC^* mice at P24 (100µg tamoxifen on P2 & P3). **(B)** Wholemount immunostaining for CD31 in cerebral cortex slices of P31 control and *Alk1^iΔAEC^* mice (75µg tamoxifen on P2).

### *Alk1^i^*^Δ^*^AEC^* mice exhibited vascular smooth muscle cell defects

Large tortuous blood vessels have been described in multiple diseases including Loeys-Dietz syndrome, Marfan’s syndrome, and Arterial Tortuosity syndrome, all of which are disorders affecting the integrity of the tunica media and vascular smooth muscles^38,39^. With this insight, we sought to examine the effects of arterial endothelial *Alk1* loss on vascular smooth muscle cell (vSMC) function. We conducted wholemount immunostaining for α-smooth muscle actin (αSMA) in the nasal mucosa of *Alk1^i^*^Δ*AEC*^ mice and found that, the *Alk1^i^*^Δ*AEC*^ mouse nasal vessels exhibited holes in arterial vSMC coverage (**Fig. 6A**). Likewise, wholemount immunostaining for αSMA in the cerebral cortex slices revealed a similar smooth muscle defect with gaps in coverage and disruption of the stereotyped circumferential vSMC alignment (**Fig. 6B**). In summary, these data reveal arterial endothelial deletion of *Alk1* compromised the proper coverage thus function of vSMCs in the nose and brain.

**Fig. 6.**
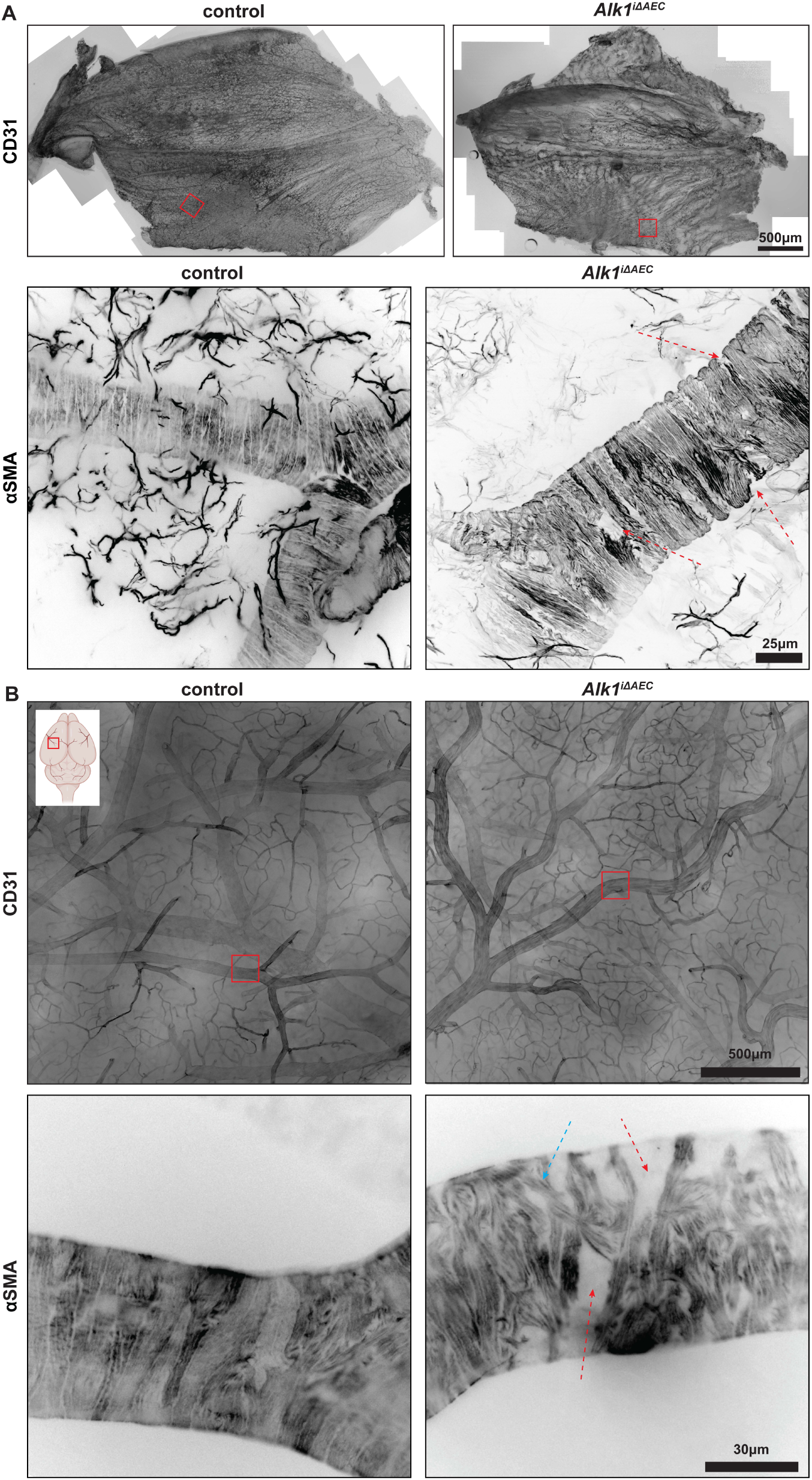
*Alk1^iΔAEC^* mice exhibited disorganized smooth muscle coverage of the nasal and cerebral vasculature. **(A)** Wholemount immunostaining for CD31 and α-smooth muscle actin (αSMA) in nasal mucosa of P103 control and *Alk1^iΔAEC^* mice (100µg tamoxifen per g body weight on P13 & P14). **(B)** Photomicrographs of wholemount immunostaining for CD31 and αSMA in cerebral cortex slices of P31 control and *Alk1^iΔAEC^* mice (75µg tamoxifen on P2). Red arrows indicate gaps in smooth muscle coverage. Blue arrow indicates regions of misaligned smooth muscle cells. Brain schematic created in BioRender.com.

### *Alk1^i^*^Δ^*^AEC^* mouse vasculature exhibited decreased expression of arterial marker *Efnb2*

Disruption of vascular smooth muscle coverage on arteries is often associated with the loss of endothelial arterial identity (reviewed in ^40^). To investigate whether *Alk1^i^*^Δ*AEC*^ mice exhibit loss of arterial identity in arteries in the nose and brain where vSMCs were disorganized, we examined the expression of an arterial endothelial marker *Efnb2*, using the *Efnb2^H2B-GFP^* reporter allele^34^. In control mice we found widespread expression of H2B-GFP in the arterial vessels; however, in *Alk1^i^*^Δ*AEC*^ mice we uncovered loss or reduction in *Efnb2^H2B-GFP^* expression in arterial branches in both the nasal mucosa (**Fig. 7A, B**) and brain and (**Fig. 7C**). The vessels exhibiting reduced *Efnb2^H2B-GFP^* expression were the arteries which had undergone Cre-mediated recombination thus loss of *Alk1*, as demonstrated by expression of the *Rosa26^Ai75^* Cre recombinase reporter (**Fig. 7B**). Together, these data indicate that arterial endothelial deletion of *Alk1* results in decreased arterial identity as demonstrated by expression of the arterial marker *Efnb2*.

**Fig. 7.**
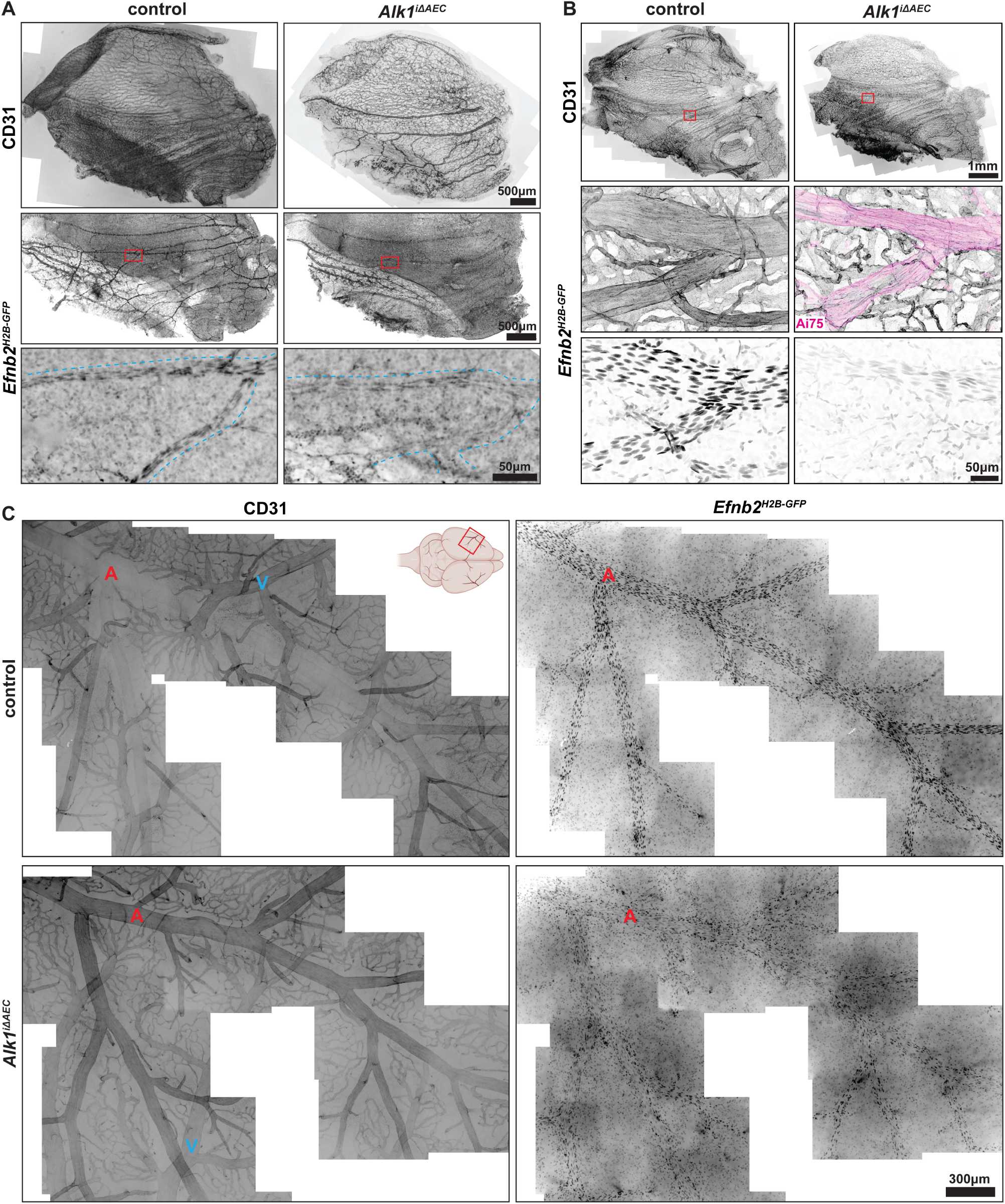
*Alk1^iΔAEC^* mice exhibited downregulation of arterial marker *Efnb2* expression in the nasal and cerebral arterial vasculature. (A) Wholemount immunostaining for CD31 and expression of *Efnb2^H2B-GFP^* arterial endothelial reporter in nasal mucosa of P13 control and *Alk1^iΔAEC^* mice (100µg tamoxifen per g body weight on P13 & P14). Red boxed indicates location of inset images. Black dashed line indicates position of artery. **(B)** Wholemount immunostaining for CD31 and expression of *Rosa26^Ai75^* Cre recombinase reporter and *Efnb2^H2B-GFP^* arterial endothelial reporter in nasal mucosa of P13 control and *Alk1^iΔAEC^* mice (25µg tamoxifen on P2). Red boxed indicates location of inset images. **(C)** Wholemount immunostaining for CD31 and expression of *Efnb2^H2B-GFP^* arterial endothelial reporter in cerebral cortex slices of P13 control and *Alk1^iΔAEC^* mice (25µg tamoxifen on P2). Mouse schematic created in BioRender.com. A and V label arteries and veins, respectively.

### *Alk1^i^*^Δ^*^AEC^* mice exhibited impaired cerebral vascular constriction and acetylcholine-induced vasodilation

Recent work from our group has identified a key role for dysregulation of arterial tone in the developmental pathophysiology of arteriovenous malformations, which are high-flow vascular anomalies that shunt blood directly from arteries to veins by displacing intervening capillaries, capable of causing both hemorrhage and ischemia^41–43^. Given the vessel enlargement and defects in vSMC organization, we hypothesized that arteries in *Alk1^i^*^Δ*AEC*^ mice exhibit defects in regulation of arterial tone, or the degree of vascular constriction and dilation. Using an in vivo assay for assessment of arterial tone^41,44^, we measured diameters of pial arteries through an open cranial window in anesthetized *Alk1^i^*^Δ*AEC*^ and control mice. We measured vessel diameters at baseline with topical administration of artificial cerebrospinal fluid (aCSF), in a vasodilated state with aCSF containing acetylcholine, and at a maximally dilated state with aCSF devoid of Ca^2+^ and containing 1 μM of the L-type Ca^2+^ channel antagonist, nifedipine. Acetylcholine-induced vasodilation is a measure of the pial artery vasodilation in response to endothelial nitric oxide synthase (eNOS) activation, whereas nifedipine maximal dilation is a measure of the arterial dilation when L-type calcium ion channels have been inhibited and vSMCs fail to contract. We found both acetylcholine-induced vasodilation and tone of pial arteries to be reduced in the *Alk1^i^*^Δ*AEC*^ mice relative to controls with the baseline dilation being significantly greater than that of controls (**Fig. 8**). Together, these results identify a key role of arterial endothelial *Alk1* in the regulation of cerebrovascular contractility and acetylcholine-induced vasodilation.

**Fig. 8.**
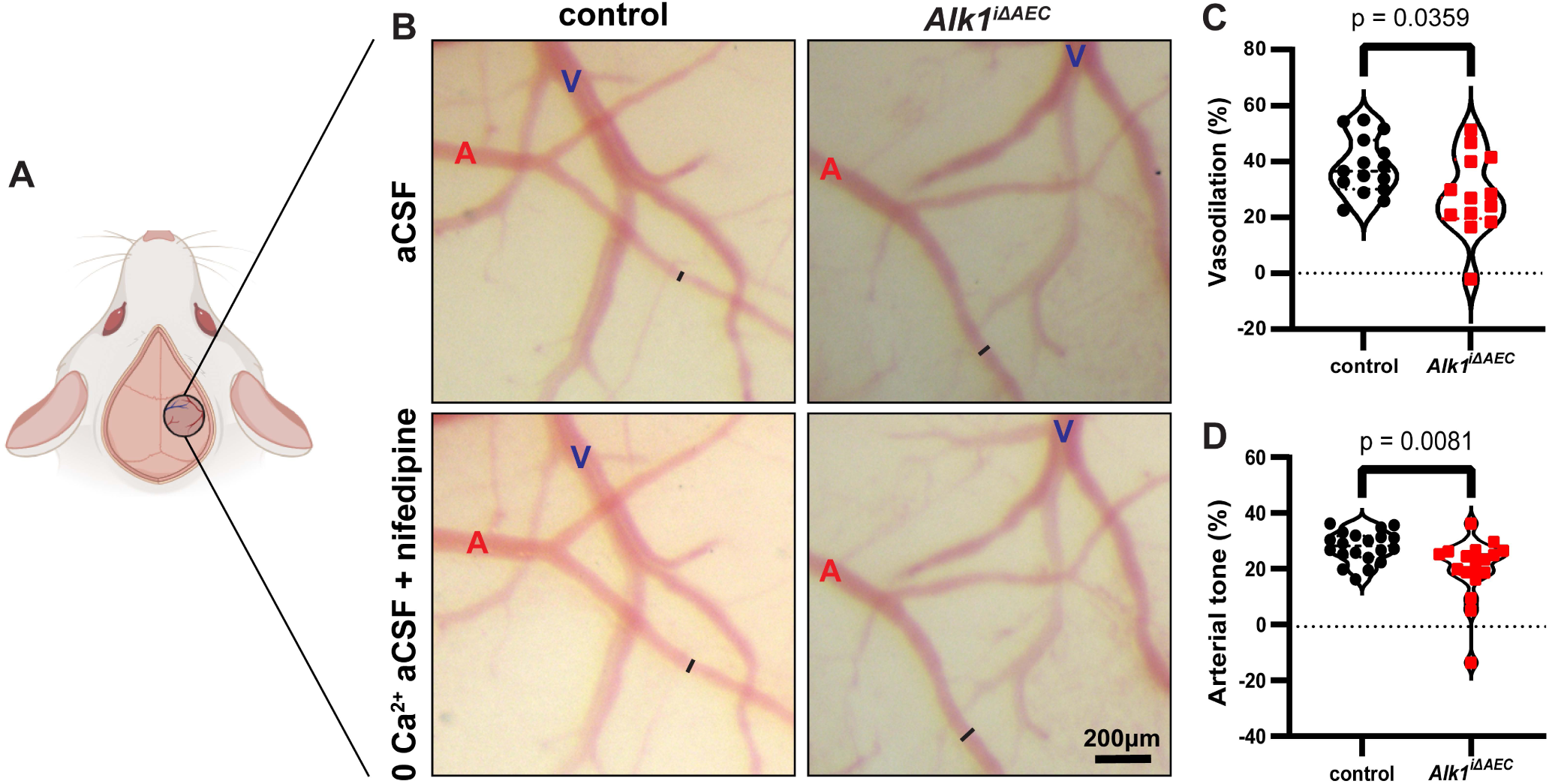
*Alk1^iΔAEC^* mice exhibited impaired in vivo cerebral vascular tone and significant changes in acetylcholine-induced vasodilation in arteries. (A) Schematic demonstrating location of pial vessel imaging during in vivo arterial tone assay. **(B)** Representative images of pial vasculature in live P13 control and *Alk1^iΔAEC^* mice (100µg tamoxifen on P2 & P3) visualized through an open cranial window. aCSF, artificial cerebrospinal fluid. **(C)** Quantification of changes in acetylcholine-induced vasodilation between control and *Alk1^iΔAEC^* mice (unpaired Student’s t-test). **(D)** Quantification of changes in arterial tone between control and *Alk1^iΔAEC^* mice (Mann-Whitney U test).

## Discussion

Here, we report that arterial endothelial *Alk1* deletion in mice elicited similar lesions as those manifested in HHT patients. We show that somatic mutation of *Alk1* in AECs results in hemorrhagic vascular malformations in the nose and brain including vessel enlargement and tortuosity, disorganization of vascular smooth muscle coverage, loss of arterial identity, and defects in vasoregulation. Taken together, these data delineate a previously unappreciated arterial contribution to HHT pathophysiology, suggesting that somatic *Alk1* deletion in AECs of mice leads to reduced arterial identity, thus disorganized vSMC coverage and defective arterial vasoregulation, which predisposes arteries to malformation and bleeding (**Fig. 9**).

**Fig. 9.**
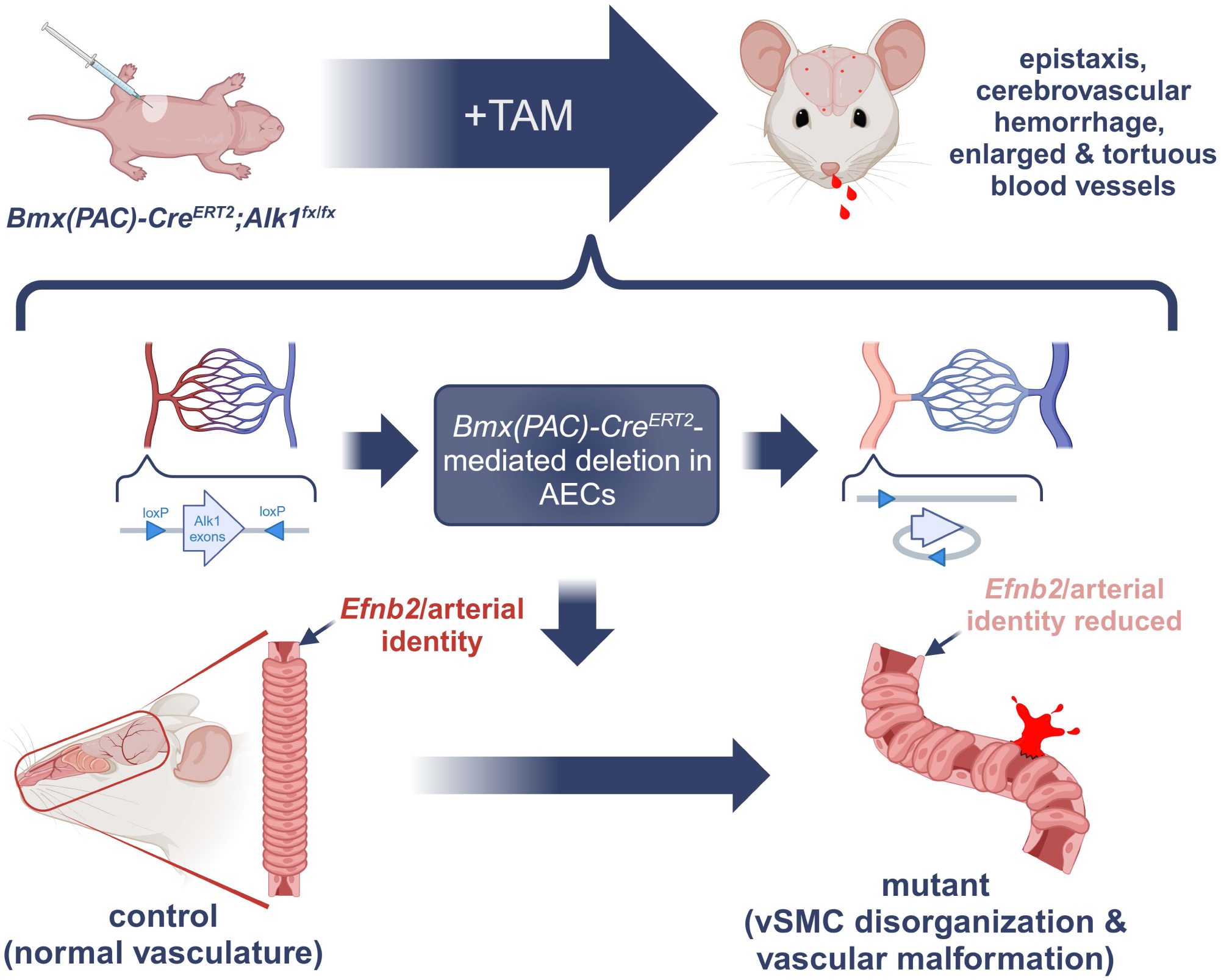
Schematic overview detailing phenotypic consequences of somatic *Alk1* mutation in arterial endothelial cells. Conceptual summary of approach and phenotypes in *Alk1^iΔAEC^* mice. AEC, arterial endothelial cell. vSMC, vascular smooth muscle cell. TAM, tamoxifen. Created in BioRender.com.

Despite being one of the most clinically significant and debilitating symptoms of HHT^9^, and many mouse mutants carrying HHT gene deletions^18,19,22,27–29,45^, epistaxis has not been reported in *Hht* mutant mice. We reported here the first mouse model of HHT-associated epistaxis. We were unable to determine the exact percentage of the mice experiencing epistaxis, given the transient nature of the bleeding episodes and the propensity of mice to clean themselves quickly. We did find about 50% of the cages containing *Alk1^i^*^Δ*AEC*^ mice showed evidence of nose bleeding which likely underestimates the true frequency of nosebleeds in these animals, as the cages are changed by animal care facility staff who do not track bleeding. The nasal vascular imaging showed 100% of vascular malformation, we believe that these mice, like patients, develop high frequency of epistaxis. Our finding presents a significant advancement in understanding pathogenesis of HHT-associated epistaxis.

The arterial endothelial specificity of the *Bmx(PAC)-Cre^ERT2^* driver is revealed by the Cre reporter assay in nasal mucosa. Unfortunately, it is technically difficulty to achieve satisfactorily signal of ALK1 immunostaining in nasal mucosa. However, single-cell transcriptomic analysis of vascular heterogeneity in the mouse nasal mucosa demonstrates that *Bmx* expression in the nasal mucosa is largely specific to the arterial endothelium^46^, supporting the utility of *Bmx(PAC)-Cre^ERT2^* to mediate arterial endothelial *Alk1* deletion in the nasal vasculature.

Arterial endothelial ALK1 protein deletion by *Bmx(PAC)-Cre^ERT2^* is shown in the cerebral cortical vasculature. Notably in the brain, multifocal microhemorrhages occurred in 72.4% of *Alk1^i^*^Δ*AEC*^ mice, without detectable symptoms of neurodysfunction. Likewise, brain AVMs in HHT patients are smaller, multifocal, and often do not require surgical intervention due to minimal symptomatic burden^35,47^. Our finding of multifocal asymptomatic hemorrhages in *Alk1^i^*^Δ*AEC*^ brains, closely mirrors human HHT manifestation.

We titrated tamoxifen dosing as an effective strategy to develop the optimal *Alk1^i^*^Δ*AEC*^ mice for modeling HHT phenotypes. While mice dosed with high doses of tamoxifen exhibited premature moribundity (median 23 days), we found that *Alk1^i^*^Δ*AEC*^ mice with a lower dose did not exhibit obviously abnormal behaviors besides spontaneous epistaxis. Importantly, *Alk1^i^*^Δ*AEC*^ mice, like HHT patients, develop compensatory cardiomegaly likely due to defects in peripheral vascular resistance. The ability to model spontaneous epistaxis without premature illness makes *Alk1^i^*^Δ*AEC*^ mice offers an ideal preclinical model for both basic discovery and therapeutic development.

The enlarged, tortuous nasal vessels in *Alk1^i^*^Δ*AEC*^ mice was revealed by MICROFIL casting. We then obtained the first immunostaining images of the nasal mucosa of mice with mutant HHT genes and also uncovered enlarged and tortuous vessels in the vasculature of the *Alk1^i^*^Δ*AEC*^ mucosa covering the nasal septum. We believe that this vascular bed in the mouse is analogous to Little’s area in humans, where the majority of epistaxis events originate^10^. These aberrant vessels in the nasal mucosa are likely the sources of severe epistaxis in *Alk1^i^*^Δ*AEC*^ mice. Both MICROFIL casting and immunostaining of the cerebral cortex of *Alk1^i^*^Δ*AEC*^ and control mice also revealed vessel tortuosity and other vascular defects accompanying the multifocal cerebral vascular hemorrhages. The defects in vascular organization of the brain were less pronounced than those of the nasal mucosa, which mirrors the fact that HHT patients develop nasal telangiectasias much more frequently than cerebrovascular AVMs^48^.

Normal function of vascular smooth muscle, is critical for vessel integrity and its disruption is associated with vascular hemorrhage^49,50^. Thinning of the tunica media in nasal vasculature of people with chronic nosebleeds has been described comprehensively by electron and light microscopy of nasal mucosal biopsies^51^, indicating that smooth muscle dysfunction may be a precursor to epistaxis. The tortuous blood vessels were accompanied by defective smooth muscle coverage characterized by loss of the compact, radial alignment in *Alk1^i^*^Δ*AEC*^ mice. This abnormal ensheathing of the artery likely predisposes this site to hemorrhage. Thus, the vSMC defect in the arteries of *Alk1^i^*^Δ*AEC*^ mice likely plays a critical role in the progression of the malformation and predisposing the vessels to hemorrhage. Endothelial *Alk1* deletion leading to smooth muscle dysregulation is supported by endothelial *L1cre* mediated *Alk1* deletion, which results in severe enlargement of the vitelline artery/vein and medial wall thinning of the vitelline artery and embryonic lethality^19^. It is well known changes from the vascular endothelium can affect vSMC differentiation and function^52,53^ and that loss of vSMC integrity is a major risk for vessel rupture^54^. Taken together these data support that endothelial ALK1 plays a critical role in regulating smooth muscle coverage, which when disrupted is a pathological mechanism for vascular malformation leading to epistaxis.

How loss of arterial endothelial ALK1 compromised smooth muscle coverage remains to be elucidated. It is possible that endothelial arterial specification is integral to vSMC coverage, thus losing arterial identity in *Alk1^i^*^Δ*AEC*^ arteries would lead to their defective vSMC coverage. Supporting this notion, we demonstrated downregulation of the arterial marker *Efnb2* in arteries of *Alk1^i^*^Δ*AEC*^ mice, accompanying dysfunctional smooth muscle coverage. Expression of *Efnb2* is a faithful marker for arterial endothelial specification^55–59^. ALK1 signaling induces expression of *Efnb2* and that loss of ALK1 is sufficient to downregulate *Efnb2* expression in mouse embryos and human cells^13,60^. Likewise, arterial specification markers *Id1* and *Hey1* are downregulated when HHT1 is deleted in the endothelium in mice ^29^. It is likely that loss of ALK1 in the arterial endothelium of *Alk1^i^*^Δ*AEC*^ mice reduces arterial programming, leading to smooth muscle defects, which predispose vessels to hemorrhage. Together, these data support a model wherein ALK1 modulation of *Efnb2* expression is a potential mechanism underlying compromised arterial vSMC coverage and hemorrhages.

Consistent with vSMC dysfunction, we identified that arterial endothelial loss of *Alk1* results in defective vasoregulation as demonstrated by reduced in vivo vascular tone and acetylcholine-induced vasodilation. These findings provide mechanistic insight into the cause of vessel enlargement and tortuosity, as the arterial contractility is disrupted. In addition, defective endothelial signaling may play a critical role in modulating vSMC functions, as increased reactive oxygen species generation is linked to compromised pulmonary artery vasomotor tone in *Alk1^+/-^* mice^61^. Given that deletion of *Alk1* reduced vascular tone in both brain and lung vasculature, it likely affects vascular tone in other tissues, including the nasal mucosa, where vascular tone measurement is not currently feasible.

We propose that defects in a feeding artery may contribute to the severe epistaxis experienced by HHT patients. This hypothesis is supported by previous work which found that there is no correlation between the size and the hemorrhage risk of a brain AVM, and that smaller AVMs with higher arterial feeding pressure are more prone to hemorrhage^62^. Our work underscores the need for clinical study of potential arterial lesions as a pathological basis in HHT-associated epistaxis.

In summary, this work demonstrates the first evidence that somatic *Alk1* loss in AECs can cause hemorrhagic vascular malformations, delineating a previously unappreciated arterial contribution to HHT pathogenesis. We believe that that this work represents a conceptual advance in our understanding of the cellular events inciting HHT-associated epistaxis, provides new avenues to better understand the molecular determinants of vascular defects leading to the devastating bleeding in patients.

## List of Supplementary Materials

Supplemental Figures 1-4

Supplemental Movie 1

Supplemental Methods

## Materials and methods

### Mice

Animals were maintained and treated in accordance with guidelines of the University of California San Francisco Institutional Animal Care and Use Committee with approved protocol AN183040. *Bmx-Cre^ERT2^* (MGI: 5513853), *ROSA26^Ai75^*(MGI: 5603432), *Alk1^fx^* (MGI: 4398904), and *Efnb2^H2B-GFP^*(MGI: 3526818) alleles have been published^19,33,34,63^. The genetic background of the mice used in this study was mixed C57BL6/J and FVB/N. Both male and female mice were used for experiments.

Tamoxifen (25, 50, 75, or 100µg; Sigma, T5648) diluted in peanut oil (Fisher Scientific, S25760) was injected intragastrically into the P2 mice unless otherwise noted. Mice injected at P13 and P14 were injected intraperitoneally with 100 µg tamoxifen per g body weight. Moribundity was determined by trained veterinary nurses in accordance with the NIH Office of Animal Care and Use guidelines. Veterinary nurses were blind to any experimental information.

### Gross pathology

Mice were anesthetized using isoflurane were exsanguinated by transcardial perfusion of PBS followed by 1% PFA. Organs, including the brain and heart, were dissected and fixed in 10% normal buffered formalin overnight at 4°C. To examine skin, fur was removed using hair removal cream according to manufacturer protocol.

### Vascular casting

Mice were anesthetized using isoflurane and were exsanguinated by transcardial perfusion of PBS followed by 1% PFA. MICROFIL:diluent:curing agent (5:4:1) was cast through the aorta. MICROFIL cast was allowed to cure at room temperature for 45 minutes and tissue was harvested, and fixed in 10% neutral buffered formalin overnight at 4°C. After serial dehydration in 25, 50, 75, and 100% ethanol at room temperature at one day intervals, tissue was then cleared in methylsalicylate before imaging. Images were captured using dissection scope and Leica LAS software.

### Fluorescent probes and immunostaining

Brains^41^ were dissected and whole-mount stained according to published protocols. Dissection and preparation of nasal mucosa is described in the Supplemental Methods. Samples were incubated in primary antibody overnight 4°C, washed three times with PBST (PBS with 0.1% Triton X-100), incubated with secondary antibody for 1hr at room temperature, again washed three times with PBST, and then mounted for imaging.

Antibodies and fluorescent probes used in this study include: (1) rat anti-CD31 (BD Pharmingen 555370; 1:500 dilution), (2) mouse anti-17-smooth muscle actin-Cy3-conjugated antibody (Sigma, C6198/F3777; 1:2000 dilution), (3) mouse anti-17-smooth muscle actin-Alexa Fluor 488-conjugated antibody (eBioscience,53-9760-82; 1:2000 dilution), (4) rat anti-VE-cadherin (BD Pharmingen, 555289, 1:200 dilution), (5) goat anti-ALK1 (R&D Systems, 1:200 dilution), (6) donkey anti-rat Alexa Fluor 488 (Invitrogen, A-11006, 1:1000 dilution), (7) donkey anti-rat Alexa Fluor 555 (Invitrogen, A21434, 1:1000 dilution), and (8) donkey anti-rat Alexa Fluor 647 (Jackson Immunoresearch Labs, 712-605-150, 1:500 dilution).

### Microscopy and imaging

Images of casted tissue and gross morphology were collected using a dissection microscope and Leica LAS software. Images of immunostained tissues were captured using either an Axioskop 2 Plus microscope (Zeiss) and Slidebook software (Intelligent Imaging Innovations) or a Yokogawa CSU-W1/SoRa spinning disk confocal on a Ti2 inverted microscope stand (Nikon) using NIS Elements software (Nikon). Multiple images of each sample were stitched together using PTGui Pro (New House Internet Services BV). Fluorescence images were adjusted for brightness and contrast using ImageJ (National Institutes of Health, USA).

### In vivo arterial tone and acetylcholine-induced vasodilation assay

Mice were injected with 100µg tamoxifen on P2 and P3. Arterial tone was assessed at P13 as described in Supplemental Materials and in previous work^41^.

### Statistics

Distributions were tested for normality using the Anderson-Darling test. Gaussian data was tested for statistical differences using a two-tailed unpaired Student’s t-test. Non-Gaussian data was tested for differences using the Mann-Whitney U-test.

## Data and materials availability

All data needed to evaluate conclusions in the paper are present in the paper and/or the Supplementary Materials. Materials may be made available by corresponding author upon reasonable request.

## Acknowledgements

We thank Dr. S. Paul Oh for providing the *Alk1^fx/+^* mouse strain and Dr. Ralf Adams for providing the *Bmx(PAC)-Cre^ERT2^* mouse strain. We thank Dr. Jennifer Hwa for initiating the cross; Joseph Maurice and Brian Jaewoo Ko for assistance with capturing brain images; Seongju Kang for arterial diameter measurements in vascular tone assay; HyoJung Heo and Cherilyn Yu for sharing breeders; and Dr. Curtis Woodford for assistance with microscopy.

