## Supplemental Figures and Methods for "Arterial endothelial deletion of hereditary hemorrhagic telangiectasia 2/*Alk1* causes epistaxis and cerebral microhemorrhage with aberrant arteries and defective smooth muscle coverage"

### Supplemental Materials

#### Supplemental Figures

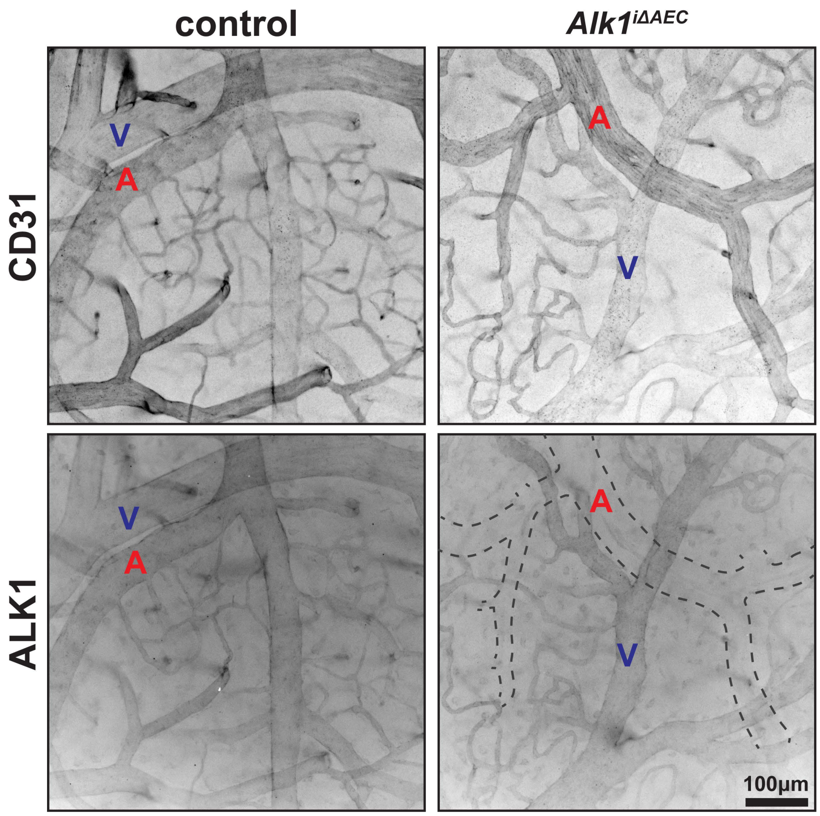


**Supplemental Fig. 1 | *Bmx(PAC)-Cre$^{ERT2}$* mediated loss of ALK1 was restricted to arterial endothelium in *Alk1$^{i\Delta AEC}$* mice induced with high tamoxifen dose.** Wholemount immunostaining for CD31 and ALK1 in cerebral cortex slices of P21 control and *Alk1$^{i\Delta AEC}$* mice (100µg tamoxifen on P2 & P3). V and A label veins and arteries, respectively.

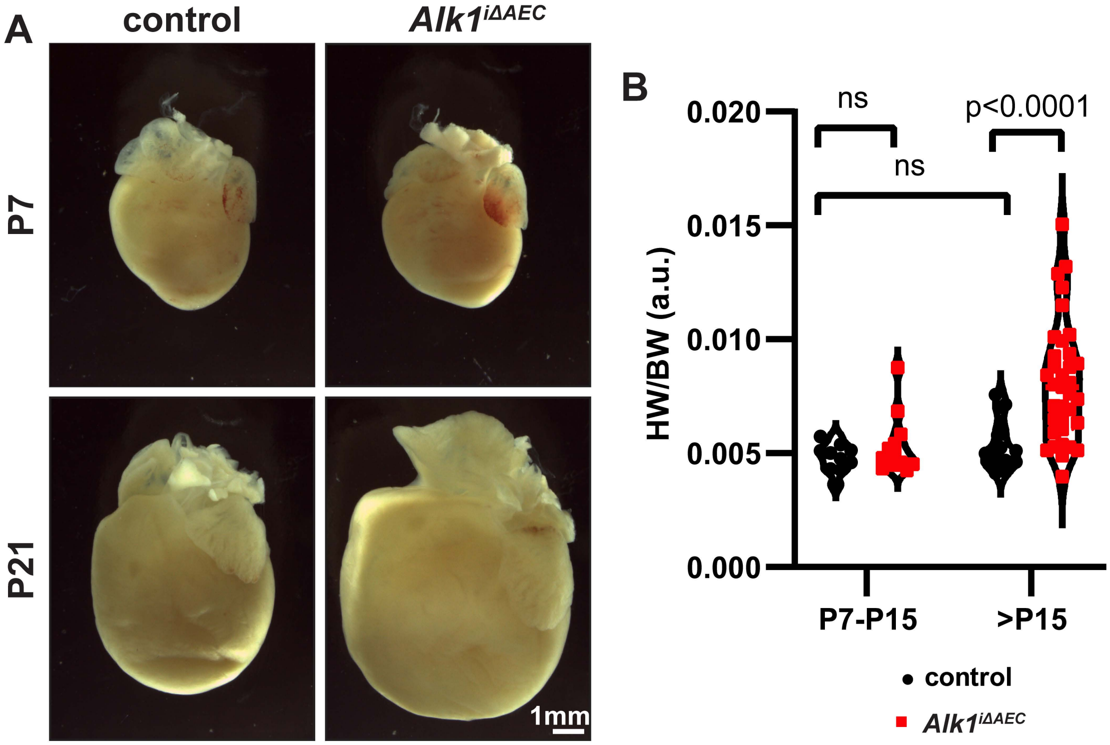


**Supplemental Fig. 2 | *Alk1$^{i\Delta AEC}$* mice developed compensatory cardiomegaly. (A)** Photographs of *Alk1$^{i\Delta AEC}$* and control (25µg tamoxifen on P2) hearts on P7 and P21. **(B)** Quantification of heart weight/ to body weight ratio (HW/BW) demonstrates a significant difference between control and *Alk1$^{i\Delta AEC}$* euthanized after P15, but not for mice before P15 (Mann-Whitney U test).

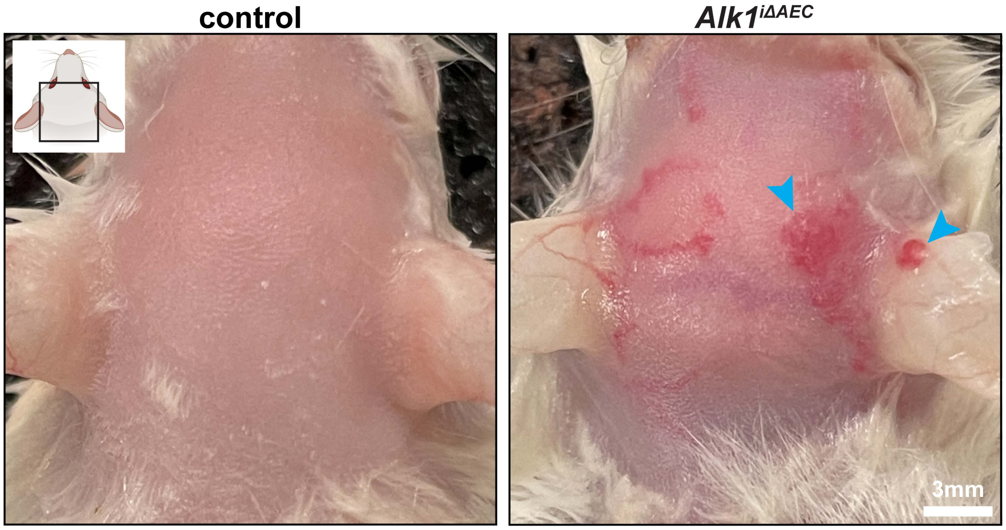


**Supplemental Fig. 3 | *Alk1*$^{i\Delta AEC}$ mice developed telangiectasia-like dermal vascular malformations.** Photographs of telangiectasia-like vascular malformations (arrowheads) present in the skin of P30 *Alk1*$^{i\Delta AEC}$ (75µg tamoxifen on P2), but not control, mice. Mouse schematic created in BioRender.com.

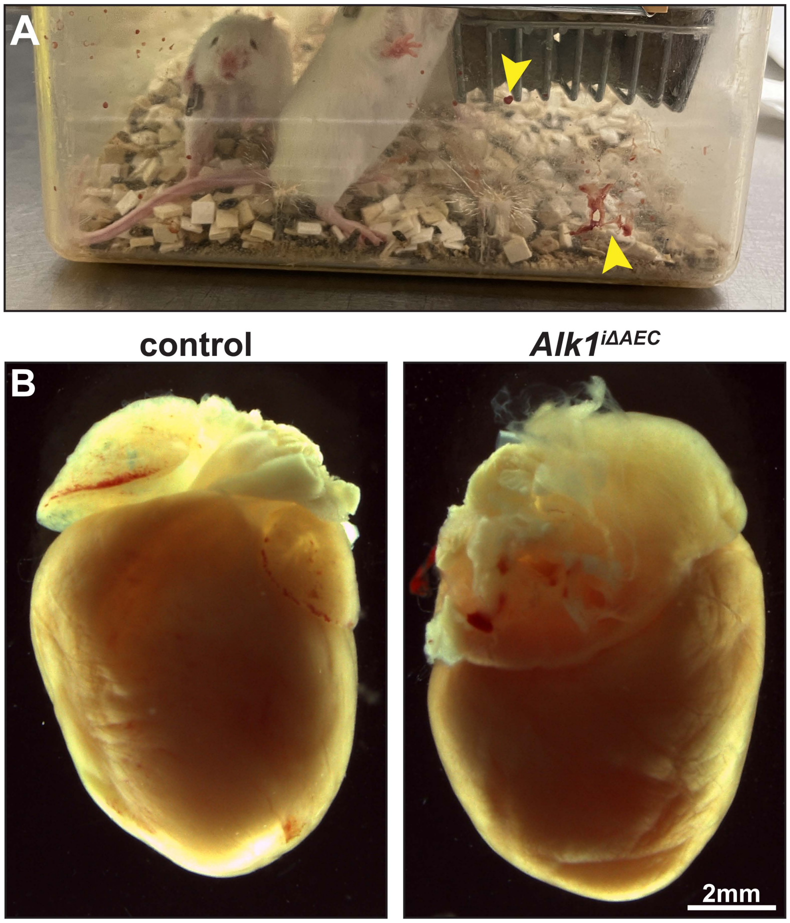


**Supplemental Fig. 4 | Induction of *Bmx(PAC)-Cre$^{ERT2}$* at P13 and P14 induced epistaxis and cardiomegaly. (A)** Photograph of blood present (arrowheads) in cage housing *Alk1$^{i\Delta AEC}$* mice. **(B)** Photographs of *Alk1$^{i\Delta AEC}$* and control hearts on P103. All mice treated with 100µg tamoxifen per g body weight on P13 & P14.

### Supplemental Movies

**Supplemental Movie 1 | Epistaxis event in *Alk1$^{i\Delta AEC}$* mouse.** Video of P33 *Alk1$^{i\Delta AEC}$* mouse (25µg tamoxifen on P2) exhibiting transient epistaxis event during handling.

### Supplemental Methods

#### In vivo arterial tone and acetylcholine induced vasodilation assessment

A surgical anesthetic plane was achieved by intraperitoneal injection (5 µl/g mouse weight) of combined 17.5% ketamine (100 mg/ml; Dechra) and 2.5% xylazine (100 mg/ml; Bimeda) diluted in physiological saline (Hospira). Surgery was conducted as described previously[1]. Briefly, a ~2mm diameter steel plate was attached to the parietal skull using a 1:1 ratio of dental cement and cyanoacrylate glue. The head plate was secured to the holding plate, and a circular craniectomy was performed. After dura opening, first artificial cerebrospinal fluid (aCSF) and then aCSF lacking calcium but containing 1 µM L-type $Ca^{2+}$ channel inhibitor nifedipine (called "$Ca^{2+}$-free aCSF + nifedipine" thereafter) were pipetted onto the exposed surface of the brain for three minutes. In some mice acetylcholine vasodilation was assessed by application of aCSF containing 10 µM acetylcholine for three minutes after the initial aCSF wash. An image was acquired and the acetylcholine was then washed out with fresh aCSF for 3 minutes. After washing, nifedipine tone was assessed described above. Images were acquired using a stereo microscope (Leica MZ12.5) with an attached camera (Zeiss AxioCam ICc5). Pial arteries were identified and analyzed using ImageJ [National Institutes of Health (NIH)]. Two locations in the arteries were analyzed per mouse. The diameters of arteries in each mouse were quantified, and results for the vascular tone and acetylcholine vasodilation in each mouse were averaged for comparison. Arterial tone was expressed as a decrease in diameter relative to the diameter obtained when incubated in $Ca^{2+}$-free aCSF + nifedipine (1 µM): [$(D2 - D1)/D2 \times 100\%$, where $D2$ = diameter in $Ca^{2+}$-free aCSF + nifedipine and $D1$ = diameter in aCSF]. Acetylcholine-induced vasodilation was expressed as the percentage change in diameter before and after ACh treatment: [$(D2 - D1)/D1 \times 100\%$, where $D2$ is diameter after ACh treatment and $D1$ is diameter before ACh treatment].

#### Preparation of nasal mucosa for immunostaining

Method was modified from published approaches[2,3]. For whole-mount preparation of the nasal mucosa, the associated skin and muscles were first stripped from the face. The skull, brain, and mandible were removed, and a blade was used to loosen the medial suture between the right and left nasal bones. Both sides of the nasal bones were gently moved and cut off at the rostral end from the underlying tissue. The anterior maxilla and frontal bone were dissected out with scissors, and the nasal mucosa along with the septum were lifted from the zygomatic bone using forceps. After removing the surrounding bones, the septum was gently gripped and removed from the nasal mucosa with fine forceps. The nasal mucosa was then fixed with 2% PFA for 2 hours at 4°C and permeabilized and blocked with blocking buffer containing 5% donkey serum in 1% Triton-X 100 in PBS overnight at 4°C. After completing immunostaining, the left and right nasal cavities were separated by cutting down the midline and then turbinate and septum mucosa were flattened and mounted for imaging.
